# Dual Transcriptional Drivers of Immune Suppression in SACC: *NOTCH1* for Cellular and *MYB* for Humoral Immunity

**DOI:** 10.1101/2025.11.17.688163

**Authors:** Guoliang Yang, Xudong Wang, Tian Ye, Tingyao Ma, Youmei Chen, Fang Nan, Qian Chen, Lu Kong, Xiaohong Chen

## Abstract

As a classic immune-“cold” tumor, salivary adenoid cystic carcinoma (SACC) represents a significant clinical challenge. Through sequencing analyses, we identify profound systemic immune dysregulation in SACC, characterized by suppressed hematopoiesis, T-cell exhaustion, compensatory myelopoiesis, and an expanded immature B-cell compartment. While lung metastases exhibit a relatively immune-engaged microenvironment enriched in tissue-resident T cells, these sites remain dominated by immature B cells. Mechanistically, this hematopoietic dysfunction originates from direct disruption of tumor-derived signaling entering the circulation, supported by detection of the SACC-signature *MYB-NFIB* fusion gene in peripheral plasma and blood cells. At the molecular level, we discover that *IL17RB/OLIG1/NOTCH1* signaling pathways engage in multicellular crosstalk via *IL33* or *TP53* to suppress T-cell activation in primary tumors. Moreover, at lung metastatic sites, the *MYB/MYBL2-NOTCH1-CXCL13-CXCR5* axis is blocked through *CD24* or *IL17RB*, thereby inhibiting B-cell maturation. Single-cell transcriptomic profiling further identifies cancer stem cells (CSCs) exhibiting dual characteristics of cell-cycle activity and immune progenitor features, a finding that clarifies the fundamental mechanism by which these cells evade immune surveillance. In conclusion, we propose that stem cell camouflage constitutes the core mechanism underlying bone marrow hematopoietic dysfunction and drives immune evasion in SACC. These findings highlight the *IL17RB-NOTCH1* signaling axis as a promising therapeutic target, offering potential to overcome the long-standing challenges associated with NOTCH1-targeted therapies in this malignancy.

**Significance:** Our novel transcriptomic analysis revealed a key G2M/CLP dual-progenitor state in SACC immunocold tumors. Tracing its origin to hematopoietic differentiation, we uncovered a *NOTCH1*-mediated dual-hit mechanism: restricting antigen presentation to silence T cells while partnering with *MYB* to block B-cell maturation. This finding pinpoints a root cause of immune evasion and establishes this pathway as a central therapeutic target.

**Graphical Abstract:** 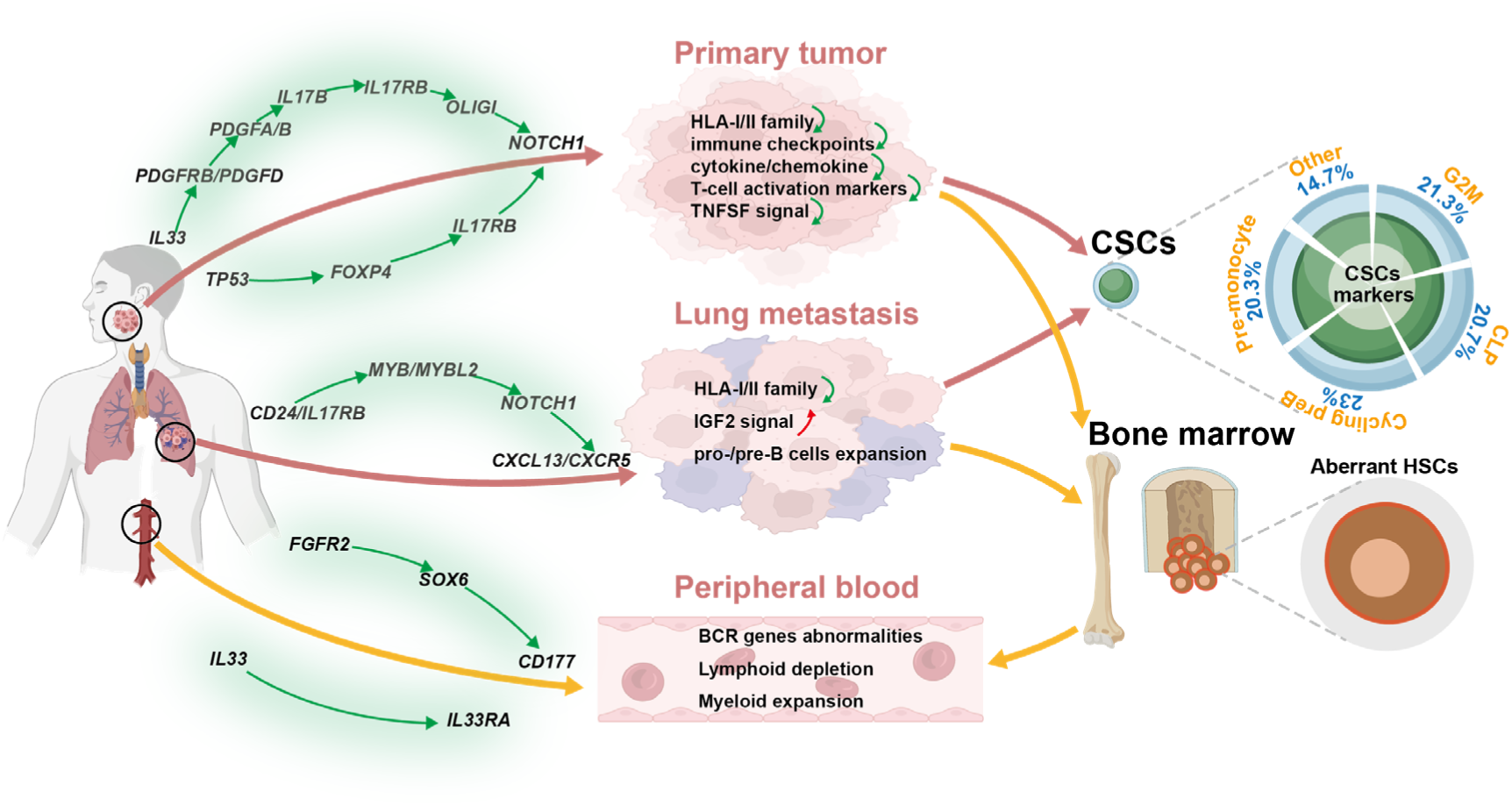

Dichotomous gene mapping and single-cell interactome reveal the microenvironmental specificity of *NOTCH1/MYB* regulation of T/B-cell differentiation and its distinct signaling flows.

## INTRODUCTION

Salivary adenoid cystic carcinoma (SACC) represents a prototypical immune “cold” tumor, exhibiting a depleted microenvironment that eludes current immunotherapies(1). Immune checkpoint inhibitors (ICIs) achieve response rates of <10% in this setting (2). Defined by *MYB-NFIB* fusions, frequent *TP53/NOTCH1* mutations, low tumor mutational burden (TMB < 2mut/Mb) and microsatellite stability, SACC follows an indolent yet ultimately lethal course: over 50% of patients develop metastatic lung disease within a decade, collapsing five-year survival from ∼70% to under 20% (3). Strikingly, our peripheral blood profiling uncovers profound co-dysregulation of B-cell receptor (BCR)-associated genes and erythroid lineage markers in SACC patients, revealing bone marrow-derived immune-hematopoietic disruption as a potential mechanism of therapeutic resistance(4–7).

*MYB-NFIB* fusions mimics a C-terminally truncated virus v-MYB, driving constitutive transcriptional activation (8). *MYB* and its homologs (*MYBL1*, *MYBL2*) play distinct roles in hematopoiesis. *MYB* is a critical transcription factor in hematopoietic stem cells (HSCs) and multilineage progenitors (9) and is involved in iNKT cell differentiation via CD1d/SLAMF/SAP signaling (10,11). In contrast, *MYBL1-* predominantly expressed in tonsillar, thymic, and splenic B cells - regulates proliferation and differentiation of mature B lymphocytes, while *MYBL2* enhances T/NK cell cytotoxicity (12). Notably, heterozygous *Mybl2*-deficient mice develop age-dependent bone marrow disorders. Tumor-expressed *NOTCH1* also exerts specific immune-modulatory effects, for example, gender-specific regulation of dendritic and T-cell-mediated anti-tumor responses in hepatocellular carcinoma (13). Although its role in T-lineage commitment (particularly γδT-cell development) is well-established, a broader understanding of how *NOTCH1* influences on immune ontogeny remains elusive(14–16). Importantly, emerging evidence reveals that tumors systemically reprogram bone marrow-derived immune cells via circulating immunosuppressive populations (17). However, the mechanisms by which SACC drivers (e.g., *MYB, MYBL1, MYBL2, NOTCH1*) manipulate hematopoietic stem cells to foster sustained immunosuppression represent a profound knowledge gap.

The Human Cell Mapping Initiative has defined 44 to 51 T-cell subtypes through transcriptomic profiling of primary lymphoid organs (bone marrow, thymus), secondary lymphoid organs (lymph nodes, spleen), and peripheral blood (18). Additionally, spatial transcriptomics now allows multi-scale immune decoding of immune architecture, mapping interactions from local tumor microenvironments to coordinated immune activity across tissues (19,20). This powerful approach has revealed distributions of epidermal/subdermal dendritic cells, tissue-resident memory T-cells (TRM), and macrophage in non-lymphoid organs, deepening insights into disease mechanisms(21–23).

We report the first detection of the *MYB-NFIB* fusion gene in a patient’s peripheral blood, employing a combined strategy of Sanger sequencing and nested PCR (Supplementary Fig S3). Furthermore, by integrating single-cell and bulk RNA-seq data from SACC tumors, metastases, and blood, together with a public single-cell atlas(24), we systematically characterized the complex immune microenvironment and regulatory networks in SACC by enrichment scores method. Through binarization and stratification of the data, we simulated gene “knockdown” and “overexpression” to reconstruct upstream regulators and downstream effectors. Notably, this integrated analysis uncovered a paradigm-shifting duality whereby oncogenic drivers not only orchestrate malignant progression within tumor cells but also, via the circulatory system, actively suppress the differentiation of immune progenitor cells, unveiling new targets for immunomodulatory therapy.

## RESULTS

### Distinct Immune Profiles in Blood, Primary Tumors, and Lung Metastases

SACC patients demonstrated systemic immunosuppression, with significantly reduced immune scores across peripheral blood (PB), primary tumors, and pulmonary metastases (Fig.1A, left), each compartment revealing a unique immune landscape.

**Fig. 1.**
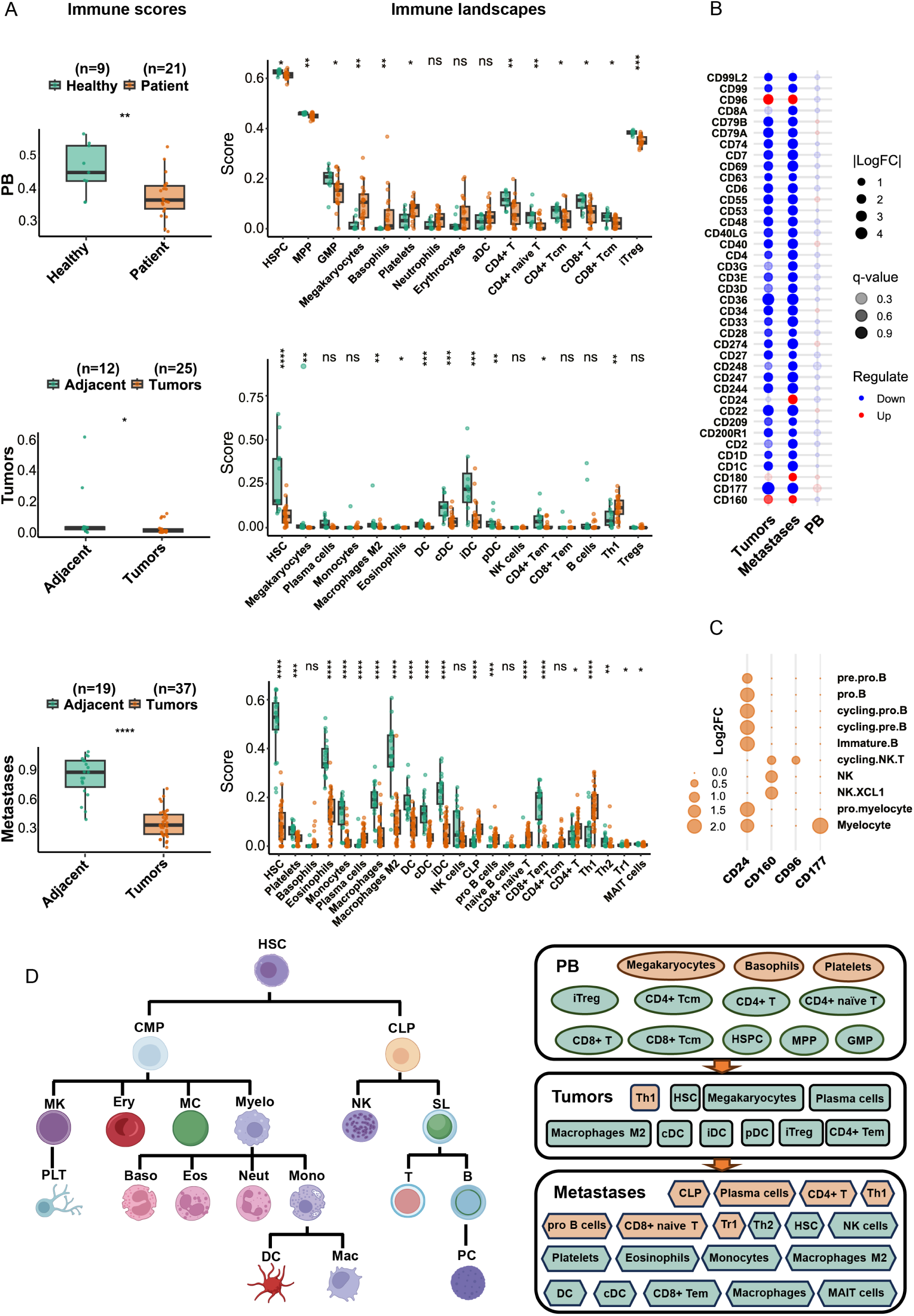
Immune profiles and LDAs expression in PB, primary tumors, and lung metastases of SACC patients. **A)** Immune scores and immune cell composition across different compartments. *Top:* PB of SACC patients vs. healthy controls; *middle:* primary tumors vs. adjacent non-tumor tissues; *bottom:* lung metastases vs. adjacent normal tissues. Light green indicates healthy individuals or adjacent non-tumor tissues, and light red indicates SACC patients or tumor tissues. **B)** Differential expression patterns of LDAs in PB, primary tumors, and lung metastases of SACC patients. The size of the bubbles represents the logFC values. **C)** Compartment-specific upregulation of LDAs defines unique immune cell signatures in PB, primary tumors, and metastatic tissues. The single-cell data were derived from the public dataset ABC. **D)** Schematic illustrating hematopoietic differentiation (Left) and multi-compartment immune cell alterations (Right). Light red and green indicate upregulation and downregulation, respectively. Mann-Whitney U test, *p<0.05 (*), p<0.01 (**), p<0.001 (***), p<0.0001 (****)*.

PB showed profound lymphoid depletion and compensatory myelopoiesis compared to healthy controls. Significantly diminished populations included hematopoietic stem/progenitor cells (HSPCs), multipotent progenitors (MPPs), granulocyte-macrophage progenitors (GMPs), CD4⁺ naïve T cells, induced regulatory T cells (iTregs), conventional CD4⁺ and CD8⁺ T cells, and central memory T cells (Tcm). In contrast, megakaryocytes, Basophils, and platelets were elevated, alongside increasing trends in monocytes, erythrocytes, and neutrophils (Fig. 1A, right, top).

Primary tumors demonstrated selective Th1 enrichment yet widespread suppression of effector lineages. Significantly reduced components encompassed HSCs, megakaryocytes, plasma cells, M2 macrophages, conventional and immature dendritic cells (cDCs, pDC and iDCs), and CD4⁺ effector memory T cells (Tem) were all significantly reduced (Fig. 1A, right, middle).

Lung metastases displayed a paradoxically’hot’ microenvironment with lymphoid activation and myeloid suppression (Fig. 1A, right, bottom). Significantly expanded populations included plasma cells, early lymphoid progenitors (CLPs and pro-B cells), and multiple T cell subsets (CD8⁺ naïve, CD4⁺, Th1, and Tr1). However, persistent deficiencies were observed in crucial cytotoxic effectors,including T cells (CD8⁺ Tem, MAIT, Th2), innate immune cells (DCs, cDCs, macrophages, M2 macrophages, monocytes, eosinophils, NK cells), and hematopoietic/prothrombotic elements (HSCs, platelets). This impairment signifies ongoing immune evasion despite local immune activation.

Leukocyte differentiation antigens (LDAs) were globally suppressed, with compartment-specific expression patterns: *CD24* and *CD180* were uniquely upregulated in lung metastases; *CD177* elevated in blood but down in both primary tumors and metastases; and *CD96* and *CD160* were elevated in both tumor compartments (Fig. 1B). Public single-cell transcriptional atlas of blood cells (ABC) (Supplemental Fig. 1A) confirms *CD24*’s expression in early B-lineage, particularly in pre-B cells, and in myeloid progenitor, *CD180* in antigen-presenting cells (e.g., B cells and DCs), and *CD96/CD160* in proliferating NK and T cells, while *CD177* marks myeloid lineages (Fig. 1C). These antigen-level shifts closely align with population changes depicted in Fig. 1D right, establishing the relationship between differentiation status and phenotypic evolution (Fig. 1D, left).

### Hematopoietic dysregulation in SACC

Blood transcriptome differentially expressed genes (DEGs) were defined using the thresholds |Log₂FC| > 1 with *p* < 0.05 (Materials and Methods). we identified 26 novel differentially expressed transcripts in PB, including 15 *IGH* genes (10V, 4D, 1J), 5 light chain genes (3 IGκ, 2 IGλ), and 6 non-antibody genes (4 non-coding RNAs, 1 pseudogene, 1 unconfirmed transcript), validated by IMGT® (25). Except *IGHV1-46, IGHD4-17, IGLV3-25,* and *LINC02593,* all showed upregulated (Fig. 2A, Supplementary Table S2). These BCR anomalies reflect stage-specific developmental failures: altered D-segment usage during the pro-B stage (*IGHD4-17* suppression with *IGHD3-10* dominance), defective VDJ recombination or pre-BCR checkpoint failure in the pre-B stage (*IGHV3-74-1* overexpression with low *IGHV1-46*), and immature B cell maturation arrest (low *IGLV3-25* with skewed κ-chain dominance via *IGKV1-5*) (26).

**Fig. 2.**
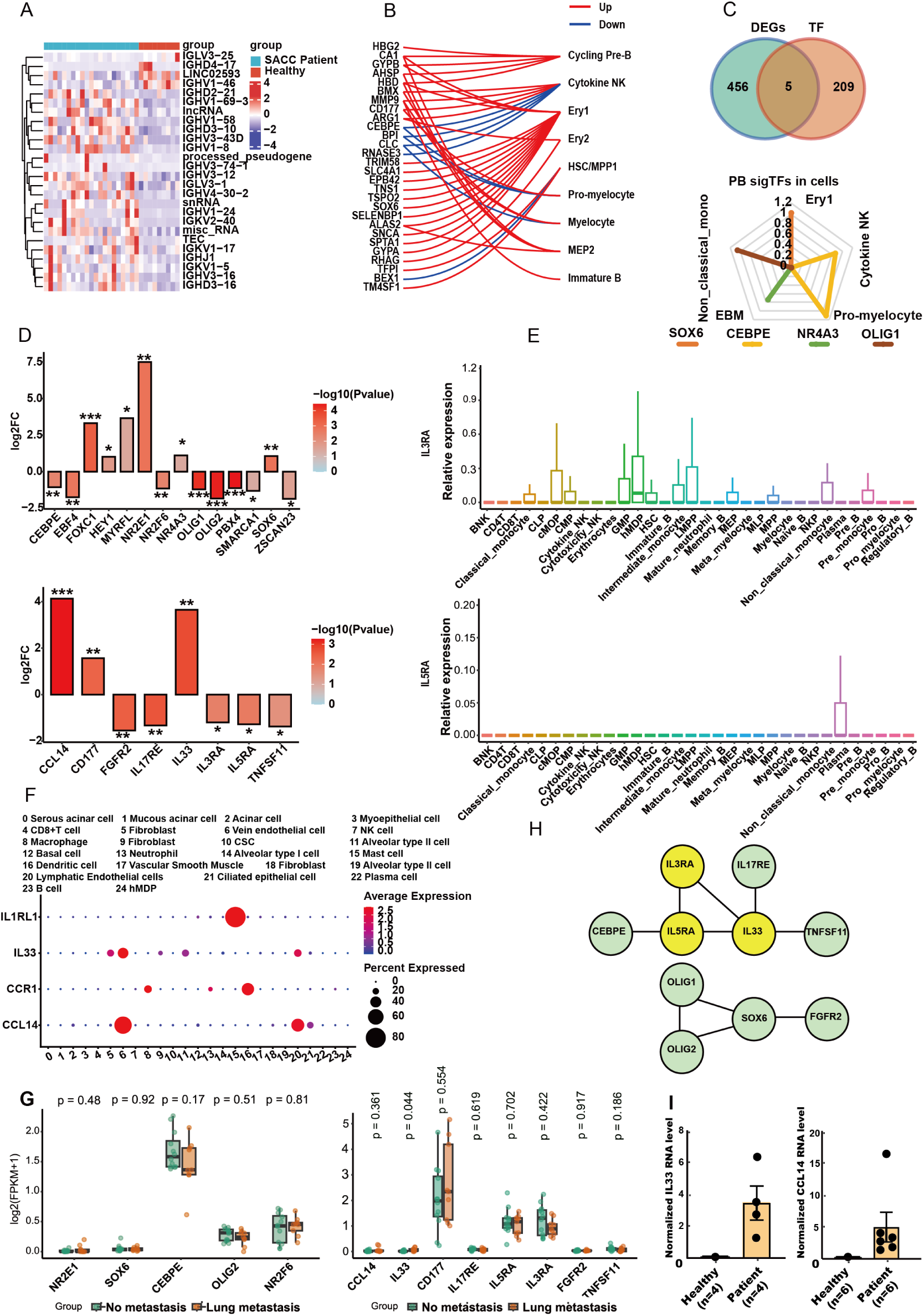
Functional Profiling of DEGs in PB transcriptomes of SACC patients. **A)** Heatmap depicting the expression patterns of 26 dysregulated novel transcripts in the peripheral blood of healthy donors (HDs) versus SACC patients. **B)** Immune cell type association of peripheral blood DEGs. DEGs identified from SACC patients compared to HDs were projected onto immune cell lineages using the ABC database. Red and blue colors indicate up-and down-regulation, respectively. **C)** Analysis of differential TFs. Venn diagram illustrating the overlap of differentially expressed TFs (upper). Radar chart visualizing the annotated immune cell types for these TFs (bottom), based on the ABC single-cell dataset. **D)** Profile of differentially expressed TFs (upper) and cytokines (bottom), shown by log_2_FC values. **E)** Expression patterns of *IL3RA* and *IL5RA* across immune cell populations, as annotated in the ABC database. **F)** Expression of *CCL14-CCR1* and *IL33-IL1R1* ligand-receptor pairs across 25 single-cell clusters from two SACC patients. **G)** Differential expression of TFs and cytokines in metastasis-free patients versus metastatic patients. Mann-Whitney U test,*p<0.05 (*), p<0.01 (**), p<0.001 (***), p<0.0001 (****)*. **H)** Protein-protein interaction network constructed using the STRING database. **I)** Validation by qPCR on blood samples confirmed elevated expression of of *IL33* and *CCL14* in SACC patients.

To further investigate this skewed differentiation, we integrated our data with the ABC database. This analysis revealed skewed differentiation toward cycling pre-B cells, immature B cells, and erythroid lineages (Fig. 2B), and further identified distinct lineage-associated transcription factors (TFs): the upregulated *SOX6* (erythroid-restricted) and *NR4A3* (immature B cells) versus the downregulated *OLIG1* (non-classical monocytes) and *CEBPE* (myeloid precursors) (Fig. 2C and D; Supplementary Table S3).

Cytokine profiling in SACC patients demonstrated dual dysregulation: elevated alarmin *IL-33* and chemokine *CCL14*, along with reduced expression of immunoregulatory receptors *IL-5RA, IL-3RA, IL-17RE, FGFR2,* and *TNFSF11* (Fig. 2D, bottom). ABC datasets confirm that *IL3RA* is broadly expressed across hematopoietic lineages, *IL5RA* mainly in plasma cells (Fig. 2E). Our single-cell mapping further showed that lymphatic/vascular endothelial-derived *CCL14* and *IL-33* interact with: (1) DCs and macrophages via *CCR1*, and (2) mast cells through *IL1R1* (Fig. 2F). Among all dysregulated factors, only *IL-33* levels specifically correlated with lung metastasis (Fig. 2G), suggesting that *IL-33* may play a pivotal role in the downregulation of *IL3RA, IL5RA, IL17RE,* and *TNFSF11*. This hypothesis supported by direct protein interaction networks (Fig 2H). Blood qPCR validated the upregulation of *IL33* and *CCL14* in SACC patients (Fig. 2I). Furthermore, Pan-cancer analysis reveals that elevated *IL33* expression is significantly associated with poor overall survival across multiple cancer types (Supplementary Fig S1).

### Cytokine Imbalance and Monocyte-Derived DC Expansion in Primary ACC

We compared cytokine profiles across peripheral blood, primary tumors and lung metastases to explore mechanisms of lymphopenia. The differentially misregulated cytokines and their ligands are detailed in Fig. 3A. Three major dysregulated cytokine networks were identified: *IL17B-IL17RB* (in both sites), *FGF9/17-FGFR1* (primary tumors), and *PDGFA/C-PDGFRA/HGFAC-MET* (metastases) (Fig. 3A). At single-cell resolution, tumor cells expressing *IL-17RB* interacted with stromal or cluster 24-derived *IL-17B* (Fig. 3D), this communication absents in circulating immune cells (Fig. 3A, bottom). Notably, IL-33 bridges PDGFRA and IL-17B–IL-17RB signaling, forming a pathogenic cascade (Fig. 3B).

**Fig. 3.**
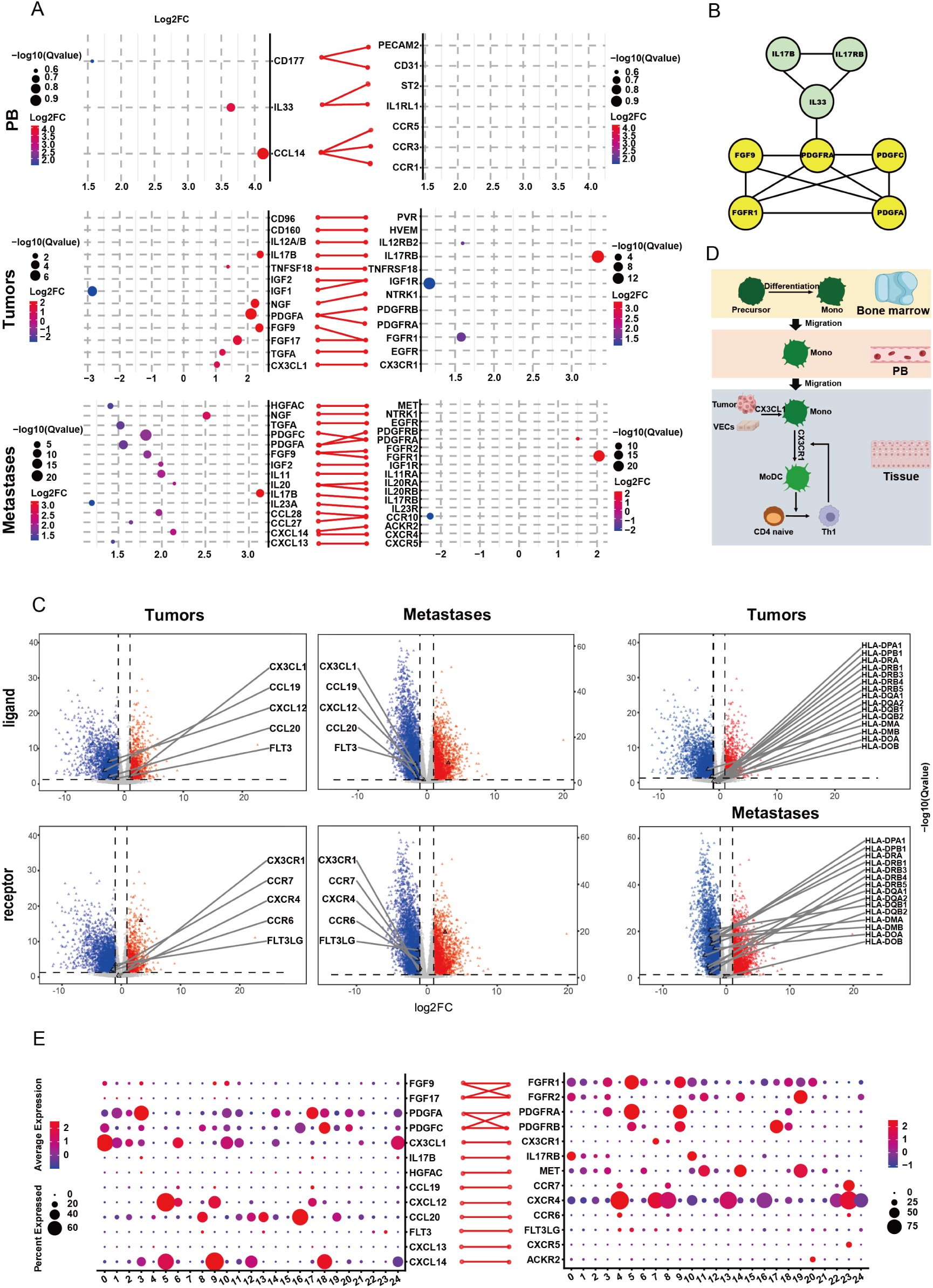
Characterization of the dysregulated cytokine network in SACC. **A)** Dot plot displaying differentially expressed receptor-ligand pairs (log_2_FC) in PB (top), primary tumors (middle), and lung metastases (bottom). The color represents the log_2_FC values. Dot size corresponds to the statistical significance (-log10 of the Q value). **B)** Predicted functional interactome among the dysregulated cytokines, mapped using the STRING database. **C)** Volcano plot highlighting key DC maturation and homing chemokines and ligands. **D)** Schematic model of monocyte differentiation into moDCs. **E)** Single-cell resolution maps from two SACC patients, annotating the specific cell types expressing dysregulated receptor-ligand pairs.

Primary tumors exhibited elevated levels of *CX3CL1*, a cytokine important for DC chemotaxis, coinciding with increased infiltration of DC precursors. Notably, the *CX3CR1* ligand (*CX3CL1*) was unchanged at primary tumors but suppressed in metastases (Fig. 3C, left and middle). Curiously, we found that key DC maturation/homing chemokines (*CCL19-CCR7, CXCL12-CXCR4, CCL20-CCR6,* and *FLT3-FLT3LG)* and major histocompatibility complex (MHC) II were downregulated across both tumor sites (Fig. 3C). Collectively, these patterns imply that tumor-infiltrating DCs are derived from monocyte precursors rather than classical myeloid progenitors.

Single-cell data further indicated that *CX3CL1* is produced mainly by tumor cells, vascular endothelial cells, and cluster 24 cells within the primary lesions, whereas *CX3CR1* is predominantly expressed on NK cells and monocytes (Fig. 3D). We propose that the *CX3CL1–CX3CR1* axis recruits and retains *CX3CR1⁺* immune cells (e.g., NK cells, monocytes), which subsequently differentiate into monocyte-derived DCs (moDCs) in response to local microenvironmental factors (Fig. 3E).

### Dysregulated Signaling in Dendritic Cell Progenitor Differentiation

Reanalysis of our single-cell data identified cluster 24 (exclusive to primary tumors) as human myeloid dendritic cell progenitors (hMDPs) (Fig 4A; Supplementary Fig S2 and Table S4) (27). Concurrent increases in blood monocyte frequency and tissue-resident Th1 cells suggest an alternative function: hMDP-derived *CX3CL1* likely recruits circulating *CX3CR1⁺* monocytes to tumor sites, where they differentiate into moDCs. These moDCs subsequently prime Th1 cell responses through antigen presentation (Fig. 3E).

**Fig. 4.**
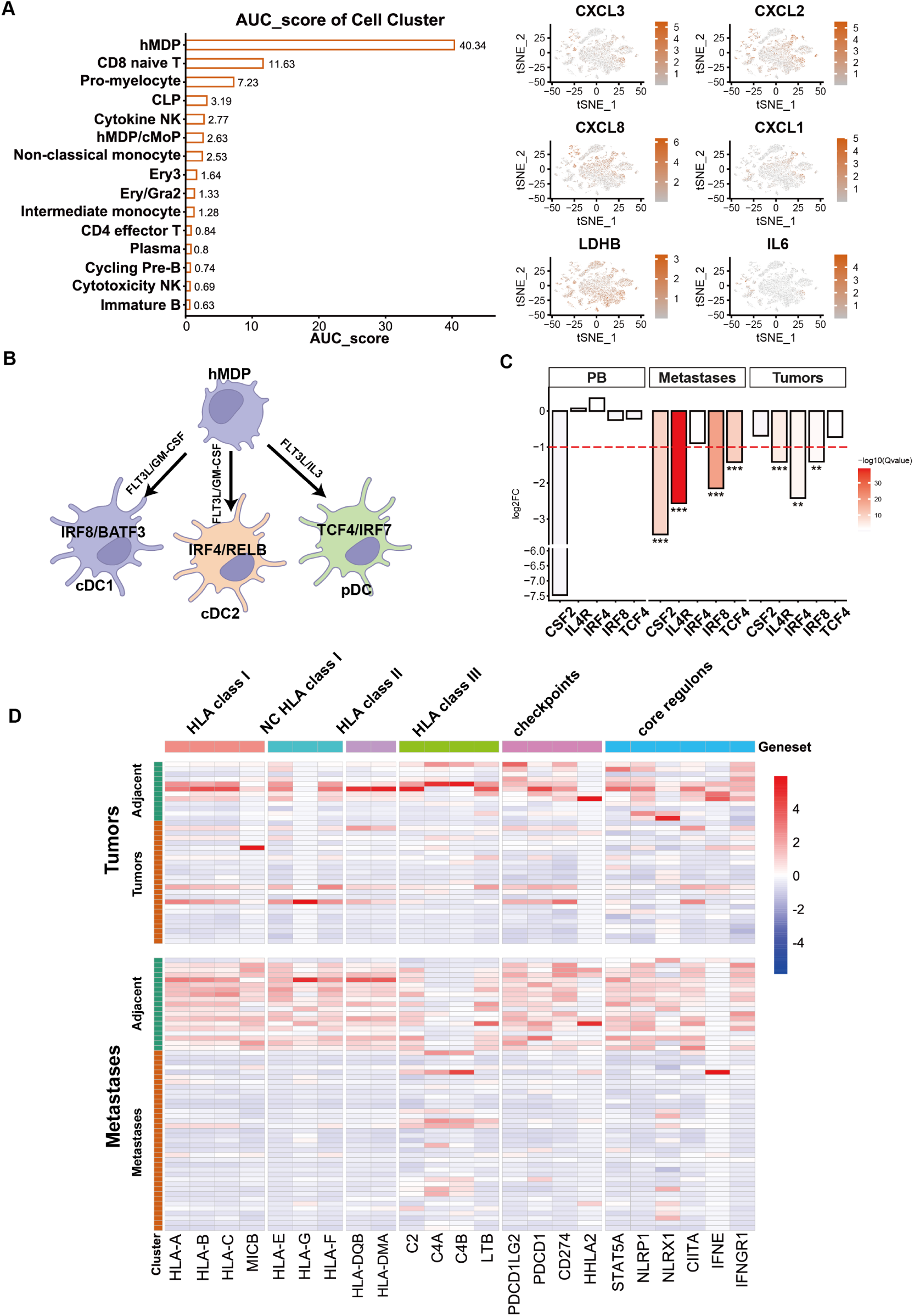
Dysregulated hMDP Differentiation into DCs Driven by the Immune Microenvironment. **A)** Cluster 24 defines an hMDP-dominant cell cluster. *Left:* AUC score (sum of AUC scores for cluster 24-specific marker genes; data from ABC). *Right:* t-SNE plot displaying cytokines common to hMDP and cluster 24. **B)** Schematic of key drivers regulating hMDP differentiation into DC subsets. **C)** hMDPs show impaired differentiation into DCs, associated with dysregulation of key drivers. Log₂FC of drivers in tumor vs. control; *|Log₂FC| > 1 and *q* < 0.05. **D)** Changes in antigen-presenting molecules across different sites. Log₂FC (tumor vs. control, |Log₂FC| > 1) is shown; red indicates upregulation; blue indicates downregulation.

Despite elevated iDC populations in metastatic sites (Fig. 1A), the absence of essential cytokines impedes their progression into functional cDCs or pDCs (Fig. 4B). Hematopoietic factor analysis revealed significant *CSF2* (GM-CSF) downregulation in metastases (trend in PB), and *IL4R* reduction in both tumor sites. Transcriptional profiling further demonstrated compartment-specific suppression of DC lineage regulators: *IRF8* (cDC1) was reduced in both tumor sites; *IRF4* (cDC2) exclusively in primary tumors; and *TCF4* (pDCs) selectively in metastases. These changes were not observed in PB (Fig. 4C).

This coincides with global downregulation of antigen-presenting machinery, including classical HLA class I (*HLA-A/B/C, MICB*), non-classical HLA class I (*HLA-E/G/F*), HLA class II (*HLA-DQB1/DMA*), HLA class III (*C2, C4A/B, LTB*), immune checkpoint (*PDCD1LG2, PDCD1, CD274, HHLA2*), core regulons (*STAT5A, NLRP1, NLRX1, CIITA, IFNE, IFNGR1*). Notably, lung metastases exhibited more pronounced suppression across *HLA classes I/II* compared to primary tumors (Fig. 4D).

### *NOTCH1* Drives an Immune-Desert Phenotype in Primary Tumors

*MYB* and *NOTCH1* overexpression define molecular features of SACC (8). To investigate their potential role in blocking DC differentiation, we employed an expression-based hierarchical strategy stratified by *MYB* or *NOTCH1* expression levels (Materials and Methods). Compared to the *MYB*-high group, the *NOTCH1*-high group displayed significant downregulation of immune-related genes (log_2_FC>1, *q*<0.05), enriching for IFN-γ response, complement inactivation, and G2M checkpoint, (Fig 5A). Moreover, *NOTCH1*-driven transcriptional effects diverged sharply between primary tumors and lung metastases (Supplementary Table S5).

**Fig. 5.**
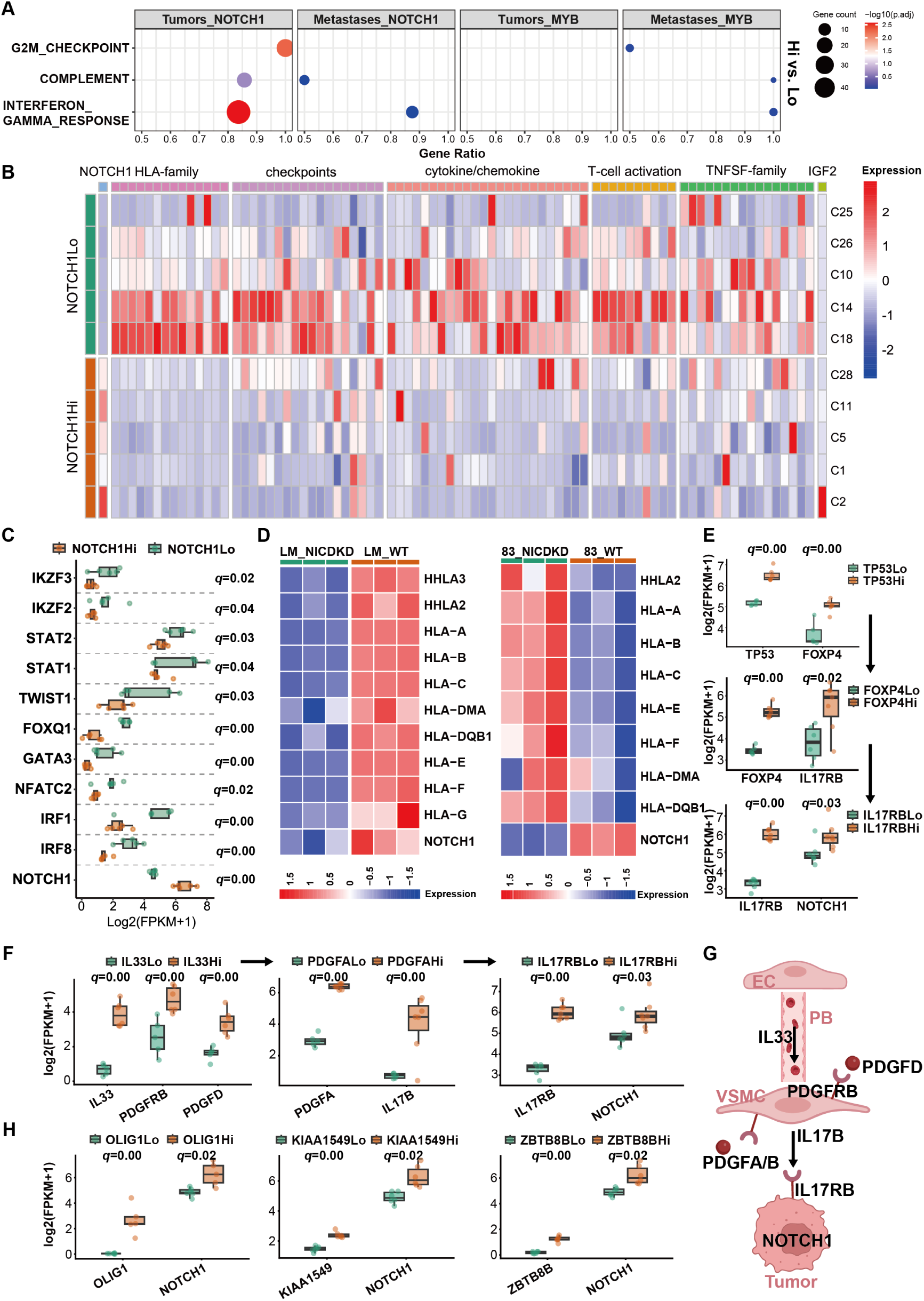
NOTCH1 drives the immunologically cold microenvironment in primary Tumors. **A)** Hallmark pathway (MSigDB) enrichment analysis of DEGs from *NOTCH1/MYB* stratification (high vs. low) in primary tumors and lung metastases (DEGs defined by DESeq2). **B)** Heatmap depicting *NOTCH1*-suppressed genes involved in antigen presentation, interferon signaling, and lymphocyte development in primary tumors. **C)** Coordinated downregulation of genes by *NOTCH1* activation across antigen presentation *(IRF8, IRF1),* immunomodulation *(GATA3, FOXQ1),* interferon response *(STAT1/2),* and lymphocyte differentiation *(IKZF2/3)*. **D)** Changes in *HLA* expression following NICD1 knockdown in SACC-83 and SACC-LM cell lines. **E)** The *TP53→FOXP4→IL17RB→NOTCH1* signaling cascade in primary tumor cells. **F)** Stratification analyze identifies the *IL33 - PDGFRB/PDGFD - PDGFA/B - IL17B - IL17RB - NOTCH1* signaling cascade **G)** Schematic: endothelial–smooth muscle paracrine signaling upregulates *NOTCH1* expression. **H)** Tumor-upregulated transcription factors enhance *NOTCH1* expression.

In primary sites, *NOTCH1*-downregulated immune genes fell into five categories: (1) HLA-family, (2) immune checkpoints (co-stimulatory/inhibitory), (3) cytokine/chemokine signaling, (4) T-cell activation markers, and (5) tumor necrosis factor superfamily (TNFSF) members involved in inflammatory regulation (Fig 5B). *IGF2* was the lone upregulated growth factor, inversely correlated with HLA class II expression (Fig 5B). Mechanistically, *NOTCH1* activation co-downregulated antigen presentation machinery *(IRF8, IRF1, STAT1, NFATC2*), immunomodulators (*GATA3, FOXQ1, TWIST1*), interferon response genes (*STAT1/2*), and lymphocyte differentiation factors (*IKZF2/3*) (Fig 5C), implying NOTCH1’s role in constraining antigen presentation, interferon signaling, and lymphocyte development. *In vitro* experiments demonstrated that NICD1 knockdown (resulting in *NOTCH1* downregulation, *q* = 0.00) in SACC-83 cells restored HLA molecule expression, confirming the conserved role of the *NOTCH1* pathway (Fig 5D). In contrast, NICD1 knockdown in the SACC-LM cell line (no significant change in *NOTCH1* expression, *q* = 0.44) instead led to downregulation of HLA molecules.

Extending this stratification approach, we identified tumor cell-intrinsic elevation of *IL17RB* expression as a direct transcriptional inducer of *NOTCH1* upregulation. Crucially, *TP53,* frequent mutations in SACC, drive constitutive *FOXP4* activation (as its direct transcriptional target), which subsequently upregulates both *IL17B* and *IL17RB* expression. This cascade establishes the mutant *TP53→FOXP4→IL17RB→NOTCH1* signaling hierarchy as the core driver of pathogenic *NOTCH1* overexpression in primary ACC (Fig 5E).

Given the high frequency of *PDGFRA* mutations in SACC fibroblasts, we investigated potential upstream regulators. Both *PDGFRA* ligands, *PDGFA* and *PDGFB* (expressed by vascular smooth muscle cells, VSMCs), directly induce *IL17B* expression in VSMCs. Reciprocally, EC-derived *IL33* stimulates *PDGFRB* and *PDGFD* expression in VSMCs, establishing a pathogenic paracrine cascade in the primary ACC microenvironment: *IL33 - PDGFRB/PDGFD - PDGFA/B - IL17B - IL17RB - NOTCH1* (Fig 5F and G). Furthermore, we identified three transcription factors, *OLIG1, KIAA1549* and *ZBTB8B*, that promote *NOTCH1* expression, with *OLIG1* is only regulated by *IL17RB* (Fig 5H).

### Impaired Humoral Immunity in Lung Metastasis via *NOTCH1*-Mediated Dysregulation of the *CXCL13-CXCR5* Axis

At metastatic sites, *IL17RB* emerged as an upstream regulator of *MYB* rather than a direct activator of *NOTCH1* (Fig 6A), consistent with our previous findings that *MYB*-driven non-canonical *NOTCH1* signaling promotes lung metastasis in ACC (28). Stratification analysis further revealed that high *NOTCH1* expression did not directly inhibit *HLA* genes or T-cell activation markers. Instead, *NOTCH1* selectively downregulated the expression of *IL22RA2, IL20RA, IGF2BP2, CD226, CD14, CXCL10, CXCL11,* and *CXCL13* while upregulating *IGF2*, a known repressor of *HLA* class II molecules (Fig. 6B). Intriguingly, *CXCL13*, which is highly expressed in metastatic lesions, potently upregulated both *HLA* class I and II expression (Fig 6B), suggesting that *NOTCH1* indirectly impairs DC function by reducing *CXCL13* and/or inducing *IGF2*-mediated *HLA* class II suppression.

**Fig. 6.**
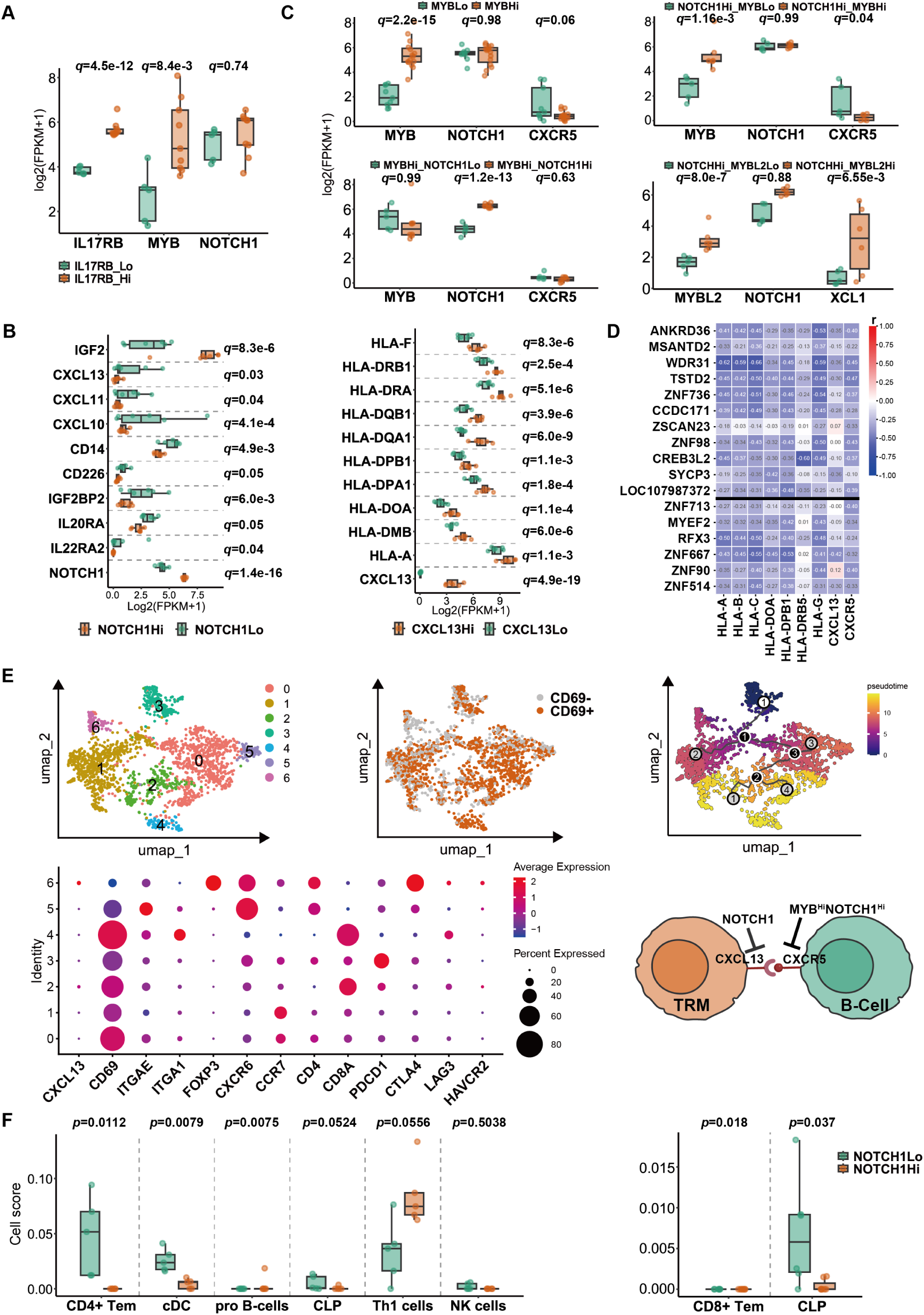
Tumor immune escape mediated by the *CXCL13-CXCR5* axis. **A)** *IL17RB* as an upstream regulator of *MYB* in metastasis. **B)** Stratification analysis of *NOTCH1-*suppressed genes (left) and *CXCL13*-mediated suppression of *HLA class I/II* (right) in lung metastases. *q*-values from DESeq2. **C)** Metastasis stratification by *MYB/MYBL2* and *NOTCH1* expression: downregulation of *CXCR5* and *XCL1* in *MYB/MYBL2*-high/*NOTCH1*-high vs *MYB/MYBL2-*low/*NOTCH1*-high groups. **D)** Negative correlation screening Identifies 17 genes (6 TFs) suppressing HLA and T cell activation. **E)** CD8⁺ T cell (cluster 4) subset analysis: transcriptionally distinct TRM populations, a CXCL13-producing Treg TRM, and a differentiation trajectory from activation to suppression (UMP and bubble plot). Schematic of TRM-primed CD8⁺ T cell/B cell interactions. **F)** *NOTCH1*-driven immune cell alterations in primary (left) and metastatic lung tissues (right).

To further dissect this axis, we stratified metastatic samples into three groups based on *MYB* and *NOTCH1* levels: (1) *MYB*-high vs. *MYB*-low, (2) *MYB*-high/*NOTCH1*-high vs. *MYB*-low/*NOTCH1*-high, and (3) *MYB*-high/*NOTCH1*-high vs. *MYB*-high/*NOTCH1*-low. In the second comparison, *CXCR5* was downregulated nearly 20-fold (Fig 6C). Furthermore, we found that high *MYBL2*/high *NOTCH1* vs low *MYBL2*/high *NOTCH1* significantly downregulated nearly 20-fold in *XCL1*, a key chemokine required for cDC1 recruitment via *XCR1* (Fig 6C). Negative correlation screening identified 17 genes that concurrently suppressed *HLA* genes and T cell activation markers (Fig 6D). The majority also downregulated *CXCR5* or *CXCL13*, implicating disruption of the *CXCL13–CXCR5* axis as a key mechanism underlying antigen presentation failure. Core transcriptional repressors driving this network included *ZNF713*, *MYEF2*, *RFX3*, *ZNF667*, *ZNF98*, and *ZNF514* (Fig 6D).

Single-cell analysis revealed *CXCL13* secretion predominantly by CD8⁺ T cells versus *CXCR5* expression confined to B cells (Fig. 3D), aligning with enrichment of CLP, pro-B, and naïve B cells in metastases (Fig. 1A). It suggested that *CXCL13-CXCR5* axis primarily governs B-cell-mediated antigen presentation at metastatic sites. Subset analysis of CD8⁺ T cells (cluster 4) revealed: subsets 0-5 universally expressed *CD69*; subset 4 exclusively expressed *ITGA1* (CD49A); subset 5 uniquely expressed *ITGAE* (CD103); subset 6 specifically expressed *FOXP3* and *CTLA4*; subsets 3,5,6 co-expressed *CXCR6* and *PDCD1*; and subset 1 expressed *CCR7.* These expression patterns are consistent with all six subsets representing tissue-resident memory T cell (TRM) populations, encompassing CD8⁺, CD4⁺, and Treg TRM subtypes, suggesting the predominant TRM within metastases (29). Notably, pseudotemporal trajectory analysis (monocle3) indicated a differentiation path from subset 3→1 and 1→0→2, revealing a transition from activation toward suppression within metastases. Treg TRM (subset 6) was identified as the primary source of *CXCL13* (Fig 6E, supplementary Table S6).

Cellular profiling further confirmed context-specific immunosuppression driven by *NOTCH1* in the tumor microenvironment. In primary tumors, *NOTCH1* overexpression correlated with reduced CD4^+^Tem and cDCs but expansion of pro B cells, while in metastases, it predominantly suppressed CD8⁺ Tem cells and CLP (Fig 6F).

### The *CD24-MYB* axis likely maintains bone marrow B cell stemness

Under physiological conditions, pro-/pre-B cells are largely restricted to the bone marrow. The detection of aberrant *IGH* gene expression in the peripheral blood of SACC patients, along with an expansion of pro-/pre-B cells in lung metastases, points to systemic disruption of hematopoiesis in these individuals. Supporting this, the *MYB-NFIB* fusion gene was identified in the plasma of one representative patient, as well as in cellular blood fractions (Supplementary Fig S3), indicating that tumor-derived components circulate systemically and may contribute to hematopoietic dysfunction. Notably, transcriptomic profiling of peripheral blood did not identify *MYB, MYBL1, MYBL2,* or *NOTCH1* as DEGs. We therefore propose that circulating tumor material may dysregulate *MYB* and *NOTCH1* signaling within bone marrow-resident hematopoietic stem cells, thereby impairing their differentiation. Direct mechanistic validation remains limited, as ethical guidelines preclude bone marrow aspiration in SACC patients, necessitating reliance on public blood and bone marrow datasets in this study.

Publicly available ABC data profile the expression of both *NOTCH1* and *MYB* across various HSPCs (HSC, MPP, CMP, GMP, LMPP, MLP, hMDP, etc.). *MYB,* however, exhibits a wider expression range, encompassing pro-B, pre-B, and immature B cells. Furthermore, *MYBL2* expression is more concentrated in pre-B and immature B cells, strongly suggesting a critical role for *MYB/MYBL2* in B-cell development. Additionally, aberrant upregulated *CD24* in metastases is a canonical B cell surface marker that is broadly expressed on pro-B, pre-B, immature, naïve, and memory B cells (Fig. 7A). qPCR validation confirmed *MYB, NOTCH1, and CD24* expression in PB (Fig 7B). Subsequent RNA sequencing demonstrated that *CD34* was significantly downregulated in SACC-LM cells upon dual knockdown of *MYB* and *NOTCH1* (Fig 7C), supporting a role for *MYB* in maintaining the progenitor B-cell state.

**Fig. 7.**
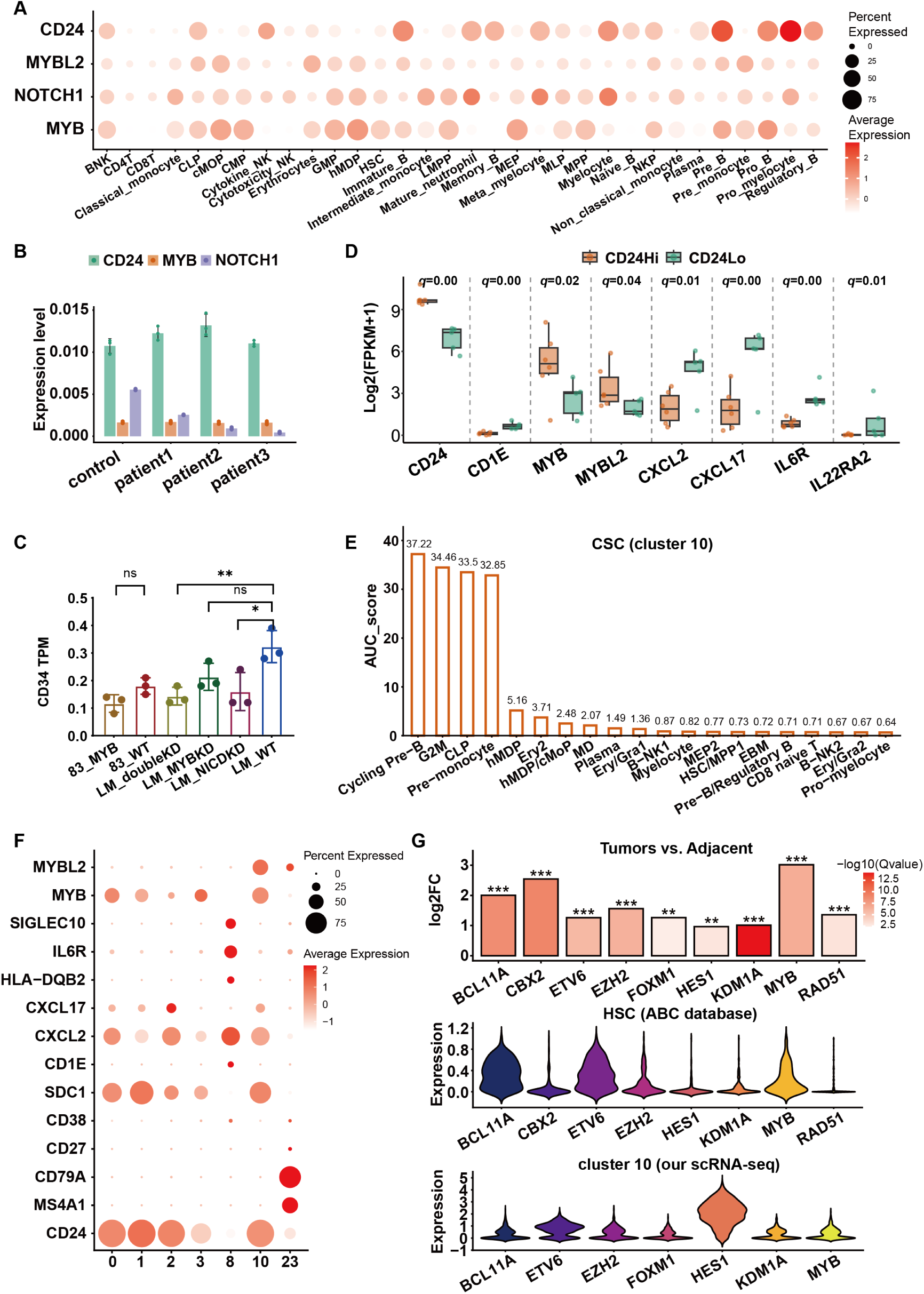
Functional evidence for ACC driver genes in bone marrow defects. **A)** Expression of *CD24, MYB, MYBL2,* and *NOTCH1* across a spectrum of 32 bone marrow and blood cell types from the ABC database. **B)** *CD24, MYB,* and *NOTCH1* are highly expressed in blood from both healthy controls and patients, as validated by qPCR. **C)** *MYB* knockdown reduces *CD34* expression in SACC-LM cells, as assessed by RNA-seq, Mann-Whitney U test,*p<0.05 (*), p<0.01 (**), p<0.001 (***), p<0.0001 (****)*. **D)** *CD24*-associated genes in metastases by stratified analysis. **E)** Based on AUC scores from the ABC database, the CSC cluster is transcriptionally most similar to cycling pre-B, G2M, CLP, and pre-monocyte cells. **F)** CSCs exhibit high *CD24* expression to engage the *SIGLEC-10*’don’t-eat-me’ signaling axis on macrophages to promote immune evasion. Clusters 0–3,10: cancer cells; 10: CSCs; 8: macrophages; 23: B cells. **G)** CSCs share a hematopoietic differentiation gene expression profile (middle and bottom), which is elevated in primary and metastatic ACC (upper).

Stratified analysis in metastases revealed that *CD24* positively regulated the expression of *MYB and MYBL2*, while negatively regulating *CD1E, CXCL2, CXCL17, HLA-DQB2, IL6R,* and *IL22RA2* (Fig 7D). These findings collectively establish that the *CD24-MYB/MYBL2* axis functions to maintain B cells in an immature, undifferentiated progenitor state.

Notably, in our single-cell sequencing data, besides *CD24*-positive B cells, CSCs (cluster 10) also consistently exhibited high *CD24* expression (Fig 7E), which promotes macrophage’don’t-eat-me’ signaling via *SIGLEC-10* (30), illustrating one mechanism by which CSCs evade immune attack.

Further analysis revealed that, unexpectedly, the marker genes defining CSC cluster shared similar transcriptional features with cycling pre-B, G2M-phase cells, CLP cells and pre-monocytes from healthy human bone marrow, as annotated in ABC databases. Signature genes of this cluster included *CD24, MYB, MYBL2* and *SDC1 (CD138)* (Fig 7F, Supplementary Table S7).

When examining the expression patterns of classical genes essential for HSC maintenance and development, we observed strong spatial concordance between *HES1* and *BCL11A* and *MYB^+^* populations, alongside epigenetic and DNA repair factors such as *EZH2, ETV6, KDM1A, RAD51,* and *CBX2*. Similarly, within the CSCs populations (cluster 10), the expression of *HES1, FOXM1, BCL11A, EZH2, ETV6,* and *KDM1A* overlapped with that of *MYB* (Fig 7F). The consistent upregulation of these genes in primary and metastatic tissues solidifies their role in SACC pathogenesis (Fig 7F). The similarity in transcriptional regulatory trajectories indicates a co-option of HSC differentiation programs by CSCs, coupled with the retention of B-cell precursor features.

## Discussion

This study demonstrates that SACC’s immunocold phenotype originates from HSPC differentiation imbalance, providing comprehensive evidence of the pathogenic hypothesis initially formulated upon detection of myeloid-derived suppressor cells (MDSCs), naïve B and double-negative T cells (DNTs) in primary tumor specimens (31).

First, the concomitant presentation of hypolymphocytosis, developmentally aberrant BCR repertoires, myeloid activation and circulating tumor-like materials in PB mechanistically substantiates hematopoietic dysregulation. Elevated plasma *IL-33* and *CCL14* further corroborate hematopoietic dysregulation, indicating non-canonical, T cell-independent type II inflammation(32). Defective hematopoietic differentiation impaired lymphoid stem cell commitment to antigen-presenting cells (APCs), evidenced by hMDP accumulation in primary tumors, downregulation of *IL3RA* (expressed in LMPP/GMP/hMDP/cMOP) and *IL5RA* (plasma cells), and broad *HLA-I/II/III* downregulation(33–35). Although monocytes differentiate into moDCs through *CX3CL1–CX3CR1* crosstalk at primary and metastatic sites, these moDCs– unlike conventional cDCs – primarily activate memory and TRM cells, driving Th1/Th17 responses(36,37). Single-cell analyses confirm resident T cells and associated Th1/Th17 gene signatures in tumor tissues (Fig. 6E). Conversely, lung metastases exhibited a significant increase in immature B cells that failed to activate resident or memory T cells via HLA class II presentation(38). This finding is supported by the presence of CD8 naive, CD4 Tcm, and Th1 cells in lung metastases. We therefore conclude that SACC immunocold primarily stems from defective APC differentiation, whereas ineffective immune inflammation in metastases results from impaired B cell–mediated antigen presentation.

Second, we show how core SACC oncogenic drivers build an immunosuppressive niche by directly and paracrinally inhibiting lymphoid progenitor cell differentiation: (1) primary site: The *IL33→PDGFRB/PDGFD→PDGFA/B→IL17B→IL17RB→OLIGI→NOTCH1* axis suppresses tumor cell *HLA-I* expression. Single-cell profiling revealed *NOTCH1* and *IL17B* expression in tumor cells and hMDPs(31). This expression pattern suggests that either *NOTCH1* signaling or *IL17B→IGF2* axis inhibits *HLA-II* expression in hMDPs, thereby impairing APC maturation and antigen presentation; (2) metastatic site: *CD24/IL17RB→MYB/MYBL2→NOTCH1→CXCL13/CXCR5* signaling restricts B cell maturation serving as a key determinant of immune exclusion. The pubic ABC dataset confirmed co-expression of *CD24, MYB,* and *MYBL2* in immature B cells, pre-B cells, or hMDP cells, while *NOTCH1* and *MYB* co-localized in progenitor cells like HSCs and LMPPs, supporting that *MYB* or *NOTCH1* abnormalities in HSCs serve as a direct cause of immune cell misdifferentiation.(39,40). Thus, *NOTCH1* constitutes the root cause of the immune desert phenotype. To date, *NOTCH1* inhibitors have not demonstrated clinical efficacy, largely owing to unacceptable toxicity profiles.

Third, Lung metastasis samples with low co-expression of *NOTCH1* and *MYB* were rarely observed(41). Furthermore, stratified analysis was limited by uniformly high *TP53* expression. Notably, *NOTCH1, MYB,* and *TP53* were all highly expressed in both tumor cells and hMDP, suggesting they may mediate lung metastasis by blocking hMDP differentiation(42). Transcriptionally, we identified *TP53* as a major upstream regulator, activating *FOXP4* to enhance *IL17RB* expression and thereby amplifying *NOTCH1* signaling. *TP53* mutations directly upregulate or disrupt *KMT6A* (*EZH2*) and *KDM5D*, contributing to chromosomal instability and tumor progression(43–45). In addition, tumor cells exploit *PDGFRA* mutations to convert *IL-33* into an oncogenic cytokine. These findings highlight *IL17RB* as a convergent therapeutic target in both primary and metastatic immune evasion programs, and preclinical efforts to target *IL17RB* are ongoing (28,46,47).

Finally, CSC clusters identified earlier exhibited marker genes closely aligned with both HSCs and progenitor B cells. This dual similarity suggests a potential role in impairing normal CLP stem cell differentiation(48). Additionally, while no viral integration was detected across five tested viruses in sequenced samples (data not shown), the origins of the *MYB* fusion and *NOTCH1* mutation remain unclear. Although somatic hypermutation (SHM) is typically confined to BCR recombination in germinal centers of lymph node, aberrant differentiation of bone marrow stem cells—which upregulates *NOTCH1* and *MYB*—may create a permissive context for SHM-like mutagenesis, warranting experimental validation(49,50).

In summary, *NOTCH1* governs T-cell/dendritic cell maturation while *MYB* drives B-cell differentiation, though precise mechanisms require validation. Future SACC therapies should target *NOTCH1/MYB*-mediated immune suppression, with stem cell immunotherapy representing a promising approach.

## MATERIALS AND METHODS

### Patient Data

This study was approved by the Ethics Committee of Beijing Tongren Hospital, Capital Medical University in accordance with the Declaration of Helsinki, with written informed consent obtained from all participants (Ethics Approval Number: TREKY2020-021). Peripheral blood samples from 22 SACC patients and 9 healthy controls were prospectively collected between 2024 and 2025. Patient characteristics are detailed in Supplementary Table S1. We additionally integrated published transcriptomic data (GSE282732) comprising: 25 primary tumor and 12 adjacent normal tissue; 37 lung metastatic tumor and 19 adjacent normal lung tissue (51). All processed with identical experimental protocols. Comprehensive patient characteristics are detailed in Supplementary Table S1, excluding those with preoperative anti-tumor therapies (e.g., radiotherapy/chemotherapy) or major immune system disorders.

### RNA Extraction and qRT-PCR

Total RNA was isolated from fresh-frozen samples using E.Z.N.A. Total RNA Kit I (Omega Bio-Tek). cDNA synthesis employed PrimeScript RT Kit (Takara) under standard cycling: 37°C/15 min → 85°C/5 sec. qPCR was performed with SYBR Premix Ex Taq (Takara) using:

95°C/30 sec initial denaturation; 40 cycles of 95°C/5 sec → 60°C/30 sec. GAPDH and actin-normalized relative expression was calculated via 2^−ΔΔCt^. Given that the Ct values of healthy controls exceeded 35 or undetected, we normalized the 2^−ΔΔCt^ values of the normal group to 0. Primer sequences are in Supplementary Table S8.

### Bulk Transcriptome Sequencing

This study utilized standardized RNA-seq protocols: Peripheral blood from 21 SACC patients and 9 healthy controls (PAXgene tubes, Agilent RIN >7) underwent Illumina NovaSeq X Plus sequencing at Xinanlou Biotech. Tissue data derived from our published cohort (GSE282732; n=89 cases) and 4 additional tumor/adjacent samples were processed identically via MGISEQ2000RS (BGI) with TRIzol extraction (RIN ≥7). All data were uniformly analyzed using HISAT2 (v2.2.1, RRID:SCR_015530) alignment to GRCh38 (ENSEMBL r112, RRID:SCR_002344), StringTie (v2.1.6, RRID:SCR_016323) quantification, and FPKM normalization. Notably, initial differential expression analysis of blood samples (n=30, ∼35,000 genes) under stringent thresholds (|Log₂FC| ≥1, adjusted *p*-value < 0.05) yielded only 13 significant DEGs. Given the dual constraints of limited sample size and high data dimensionality, we implemented a discovery-phase approach using relaxed statistical criteria (|Log₂FC| > 1, *p*-value < 0.05) to mitigate excessive filtering of biological signals. This strategy yielded 100 candidate DEGs. Critically, qPCR validation confirmed concordant expression trends for selected genes between sequencing data and experimental results.

### Pan-Cancer Analysis and Survival

Using TCGA (RRID:SCR_003193) data, we assessed *IL33* differential expression across 33 human cancers. Statistical significance (Wilcoxon test) is denoted: *\*p<0.05; **p<0.01; ***p<0.001*. Overall survival (OS) was evaluated by Kaplan-Meier analysis with time-to-event in months, implemented via R (R version 4.4.3, RRID:SCR_001905) packages survival (RRID:SCR_021137) and survminer (RRID:SCR_021094).

### Single-Cell Transcriptomics Analysis

scRNA-seq data from our published cohort (GSE216852) were reanalyzed, including one primary SACC and one lung metastasis case (3’-end sequencing) (28). Peripheral blood data integrated public repositories (24). All data were processed through a unified Seurat workflow: quality control (mitochondrial content <20%), log-normalization, PCA-based dimensionality reduction, graph-based clustering (resolution=0.5). Cluster 4 was partitioned into 7 distinct subsets after standardization (resolution=0.3), followed by pseudotime analysis using monocle3 package (RRID:SCR_018685).

### Gene Knockdown and Sequencing *in vitro*

As reported previously(28), stable *NOTCH1*-KD and *NOTCH1/MYB* double-KD cell lines were generated in SACC-LM/SACC-83 cells (RRID:CVCL_H590, RRID:CVCL_H589) using lentiviral shRNAs in pLKO.1-Puro or pCDHO-Neo-CMV-3-Flag vectors, with selection using 2 μg/mL puromycin or 1 μg/mL neomycin, respectively. Bulk RNA-seq data from these established lines are publicly available under GEO accession GSE216852. In the present study, additional RNA-seq data from NOTCH1-KD SACC-83 cell lines were newly deposited, generated using the same experimental and analytical procedures as previously described (28).

### Immune Cell Definition and Quantification

We profiled immune cell enrichment using RNA-seq data from three biological compartments: PB, primary tumors, and lung metastases. A curated panel of 75 immune cell subtypes spanning seven lineages (progenitor cells, myeloid cells, Erythroid-Megakaryocyte progenitors, B cells, CD8⁺ T cells, CD4⁺ T cells, NK cells) was analyzed using integrated marker genes from ImmuCell AI (https://guolab.wchscu.cn/ImmuCellAI, RRID:SCR_027645), XCell (http://xcell.ucsf.edu/, RRID:SCR_026446), and UCSC (https://cells.ucsc.edu, RRID:SCR_005780) databases (Supplementary Table S9).

Single-sample gene set enrichment analysis (ssGSEA, RRID:SCR_026610) was implemented via a custom Python 3.9 (RRID: SCR_008394) pipeline to quantify cell type abundance in individual samples. This method ranks genes by expression magnitude and computes a weighted cumulative enrichment score for predefined gene sets, prioritizing highly expressed markers using an exponential decay weighting factor (α = 0.25)(26). Scores were normalized by the maximum absolute deviation across gene rankings to ensure cross-sample comparability. Expression matrices underwent log₂ (FPKM + 1) transformation prior to analysis, with cell types exhibiting <10 overlapping marker genes excluded. Differential abundance between tumor samples and normal controls was assessed independently per cell type and biological compartment using the non-parametric Mann-Whitney *U* test (two-tailed; α = 0.05) after confirming non-normally distributed scores (Shapiro-Wilk test, *p*<0.05). P-values were adjusted for multiple comparisons across 75 cell types and three compartments via Benjamini-Hochberg FDR correction (significance threshold: *q*< 0.05). Critically, differential cell types were defined only when demonstrating significant enrichment (*p*<0.05) in both XCell and ImmuCellAI databases concurrently with consensus positivity in our in-house enrichment analysis.

### RNA-seq Data Stratification and Correlation in Python

The stratification and the construction of the regulatory networks were implemented in Python 3.10 using a custom computational framework (Supplementary Table S10). Gene expression values were stratified into high/medium/low tiers based on expression quantiles, with differential analysis between high and low tiers performed using the core logical framework of DESeq2 (RRID:SCR_015687). Regulatory networks were constructed via GENIE3 (RRID:SCR_000217) to predict upstream/downstream molecular interactions, followed by report generation of hub genes or specific genes (code and dependent packages can be found in GitHub, https://github.com/acc837/stratified-difference-analysis, RRID:SCR_002630). For transcription factor co-expression profiling, Spearman correlation coefficients (ρ) were computed between each transcription factor and a target oncogene across matched tumor and adjacent tissue datasets. Differential correlation analysis was conducted using Fisher’s z-transformation with Benjamini-Hochberg FDR correction (significance threshold: *q*<0.05). Tumor-enhanced transcriptional regulators were identified based on: 1) statistically significant difference in malignant versus normal tissue (FDR-adjusted *p*<0.05), and 2) absolute correlation magnitude exceeding normal tissue levels (|𝜌𝜌_𝑡umor_| > |𝜌𝜌_𝑎djacent_|). Prioritized candidates were ranked by minimal FDR and maximal absolute correlation difference (|Δρ|). Identical analytical methodology was extended to additional key pathway components.

### Assay for the *MYB-NFIB* fusion gene in blood

Tumor RNA was extracted from fresh tissue with the TRIzol Reagent (15596018, Invitrogen, Burlington, USA). Libraries were constructed using a NEBNext UltraTM II RNA Library Prep Kit (#E7775, NEB, MA, USA), and targeted region were captured using a customized Agilent SureSelectXT RNA panel (277 gene). High throughput sequencing was performed on the MGI DNBSEQ-T7 Genetic Sequencer (RRID:SCR_024847). The sequenced reads were mapped to the hg19 by Burrows-Wheeler Aligner (BWA version 0.7.15, default parameters, BWA-MEM algorithm). Gene fusion analysis was performed using STAR-Fusion (RRID:SCR_025853, version 1.8.1, default parameters) and gene fusions were validated with Integrative Genomics Viewer (RRID:SCR_011793, version 2.8.0). Plasma RNA was extracted with Starvio cfRNA Kit (N11005, Starbio, Shanghai, China) and blood cell RNA extraction was performed using RNA Extraction Kit (IVD4144, Magen, Guangzhou, China) according to the manufacturer’s instructions. Nested PCR primers were designed to span the gene fusion junction, with the outer and inner primer pairs (Supplementary Table S8). Fusion transcripts were amplified by two-round nested-PCR using 2 × Hieff® PCR Master Mix Kit (10160ES08, YEASEN, Shanghai, China) and Shanghai Hongshi Medical Technology SLAN-96S Real-Time PCR System (RRID:SCR_027087). Cycling conditions were: initial denaturation at 98°C 3 min, followed by 35 cycles of denaturation at 98°C 10 s, annealing at 60°C 30 s, extension at 72°C 30s, and a final extension at 72°C 10 min. Purified nested-PCR amplicons were subjected to Sanger sequencing using the ABI 3500Dx Genetic Analyzer (Applied Biosystems, Foster City, CA, USA) following standard protocols. Sequencing reads were analyzed using Chromas (RRID:SCR_000598).

### Statistical analysis and visualization

Differential gene expression analysis was performed using the DESeq2 package via the Dr. Tom multi-omics data mining system (https://biosys.bgi.com, Beijing Genomics Institute, RRID:SCR_027646). Intergroup comparisons were conducted using the non-parametric Mann-Whitney *U* test. All gene expression heatmaps, dot plots, box plots, violin plots, t-SNE projections, and UMAP visualizations were generated using R version 4.4.2 (with ggplot2, RRID:SCR_014601, and Seurat packages, RRID:SCR_016341). Gene expression plots for qPCR validation were created using GraphPad Prism software suite (version 10.1.2, RRID:SCR_002798). Statistical significance for DESeq2 analyses were defined as follows: *q<0.05 (*), q<0.01 (**), q<0.001 (***), q<0.0001 (****)*. ALL schematic elements were obtained from BioRender (RRID:SCR_018361).

## Author contributions

Guoliang Yang: Conceptualization, data collection, data curation and analysis, experimental validation, manuscript review, and editing; preparation of Figures 4–7, and graphic abstract. Xudong Wang: Data collection, preparation of Figures 1–3, manuscript review, and editing. Tian Ye: Algorithm development, figure preparation, manuscript review, and editing. Tingyao Ma: Data collection. Youmei Chen: Data collection. Fang Nan: Data collection. Qian Chen: Data analysis, manuscript review, and editing. Lu Kong: Conceptualization, data analysis, manuscript writing, review, and editing. Xiaohong Chen: Conceptualization, data analysis, manuscript writing, review, and editing.

## Acknowledgments

This work was supported by National Natural Science Foundation of China (grant no. 82372967). Additionally, we acknowledge AI tools (DeepSeek R1) for language refinement and hierarchical data programming.

## Data Availability

The sequencing data (including PB, tissues and cell lines) are currently being uploaded to the GSA database, and the accession number will be provided once available. Upon publication, all data supporting the findings of this study will be available from the corresponding author upon reasonable request.

## Abbreviations

ABC: Atlas of human blood cells
APCs: Antigen-presenting cells Baso: Basophil
BCR: B-cell receptor
cDCs: Conventional dendritic cells
CLP: Common lymphoid progenitor
CMP: Common myeloid progenitor
CSCs: Cancer stem cells
DC: Dendritic cell
DEGs: Differentially expressed genes
DNTs: double-negative T cells
Eos: Eosinophil Ery: Erythrocyte
GMPs: Granulocyte-macrophage progenitors
HLA: Human leukocyte antigen
hMDP: Human myeloid dendritic progenitor cells
HSC: Hematopoietic stem cell
HSPCs: Hematopoietic stem/progenitor cells
ICI: Immune checkpoint inhibitors
iDCs: Immature dendritic cells iTregs: Induced regulatory T cells
LDAs: Leukocyte differentiation antigens
Mac: Macrophage
MAIT: Mucosal-associated invariant T cells
MB: Myeloblast
MC: Mast cell
MDSCs: myeloid-derived suppressor cells
MHC: Major histocompatibility complex
MK: Megakaryocyte
moDCs: Monocyte-derived dendritic cells Mono: Monocyte
MPPs: Multipotent progenitors Neut: Neutrophil
PB: Peripheral blood PC: Plasma cells
pDCs: Plasmacytoid dendritic cells
PLT: platelet
SACC: Salivary adenoid cystic carcinoma
SHM: Somatic hypermutation
SL: Small lymphocyte
Tcm: Central memory T cells
TMB: Tumor mutation burden
TRM: Tissue-resident memory T-cells
VECs: Vascular endothelial cells

