## supplemental figure S1-S3 for "Dual Transcriptional Drivers of Immune Suppression in SACC: *NOTCH1* for Cellular and *MYB* for Humoral Immunity"

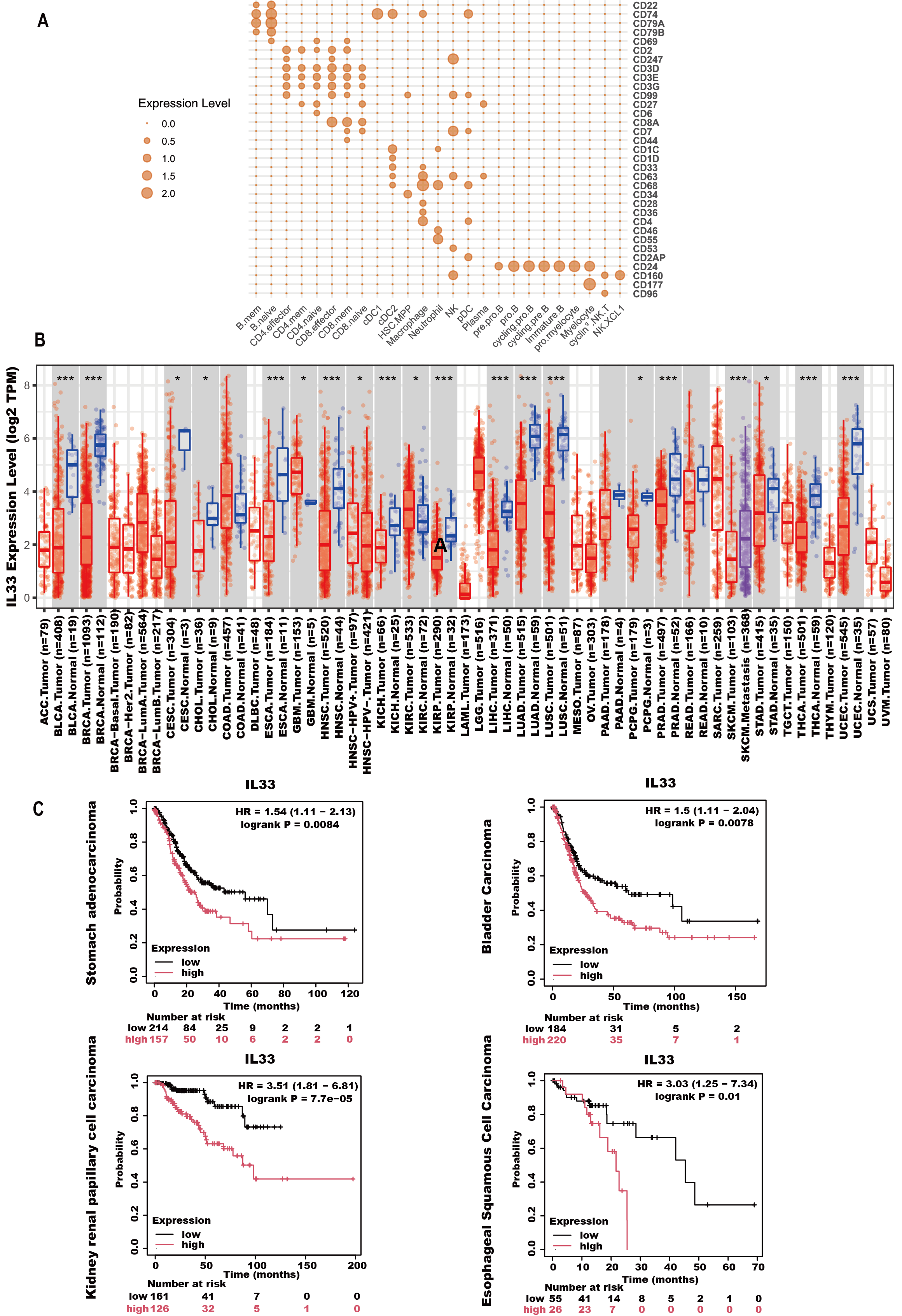

**Supplementary Fig S1. Validation of LDA and IL33 expression and their association with clinical prognosis based on public databases. A)** Differentially expressed LDAs in cancer tissues were mapped to their source immune cell types via reference-guided annotation of blood scRNA-seq ABC profiles. **B)** Pan-cancer analysis of *IL33* based on TCGA database. Blue indicates normal tissues, and red indicates tumor tissues. **C)** Association between high *IL33* expression and poor overall survival across multiple human tumors.

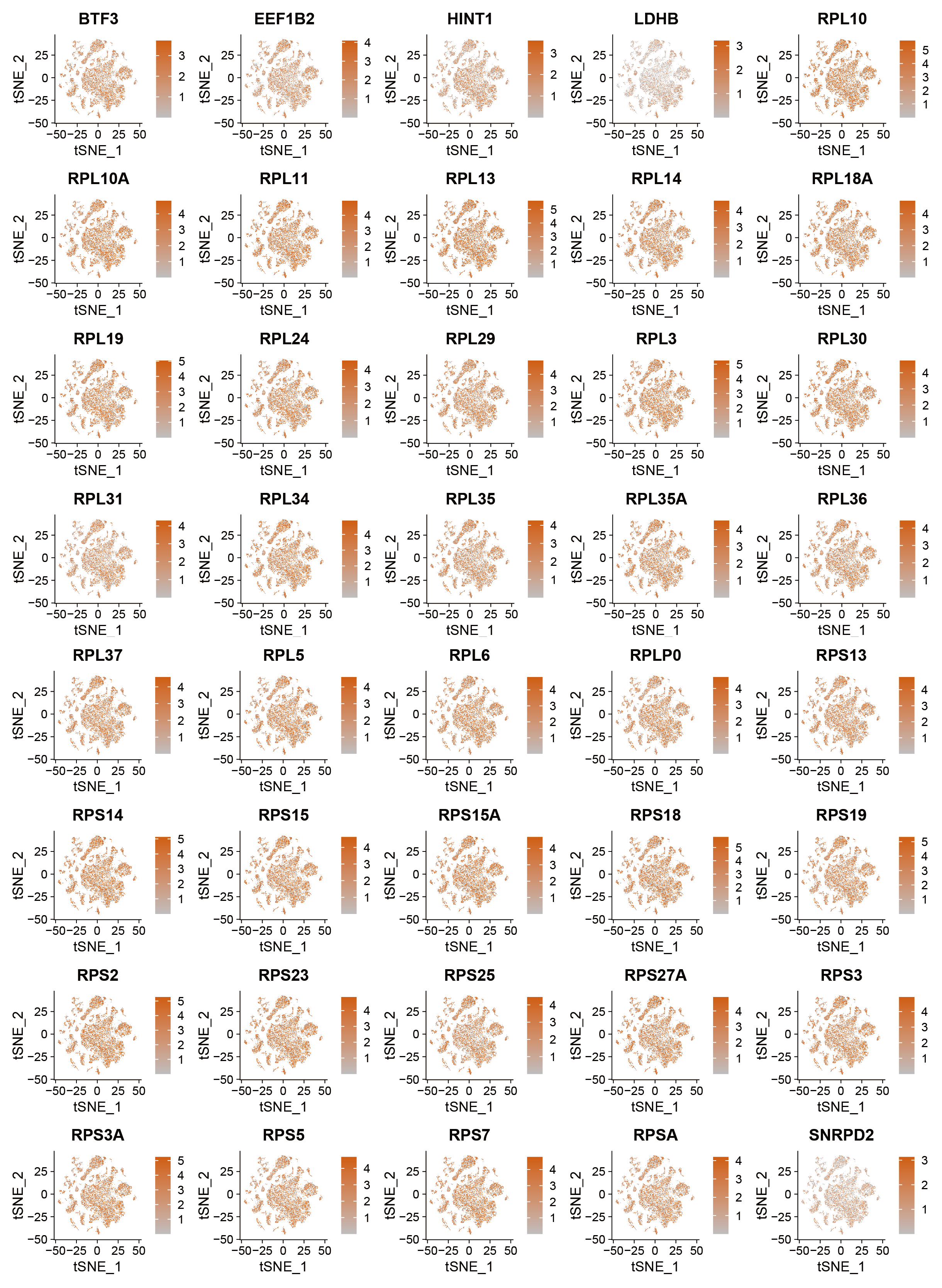

**Supplementary Fig S2. t-SNE plot visualizing the expression patterns of the top 50 marker genes for hMDP.**

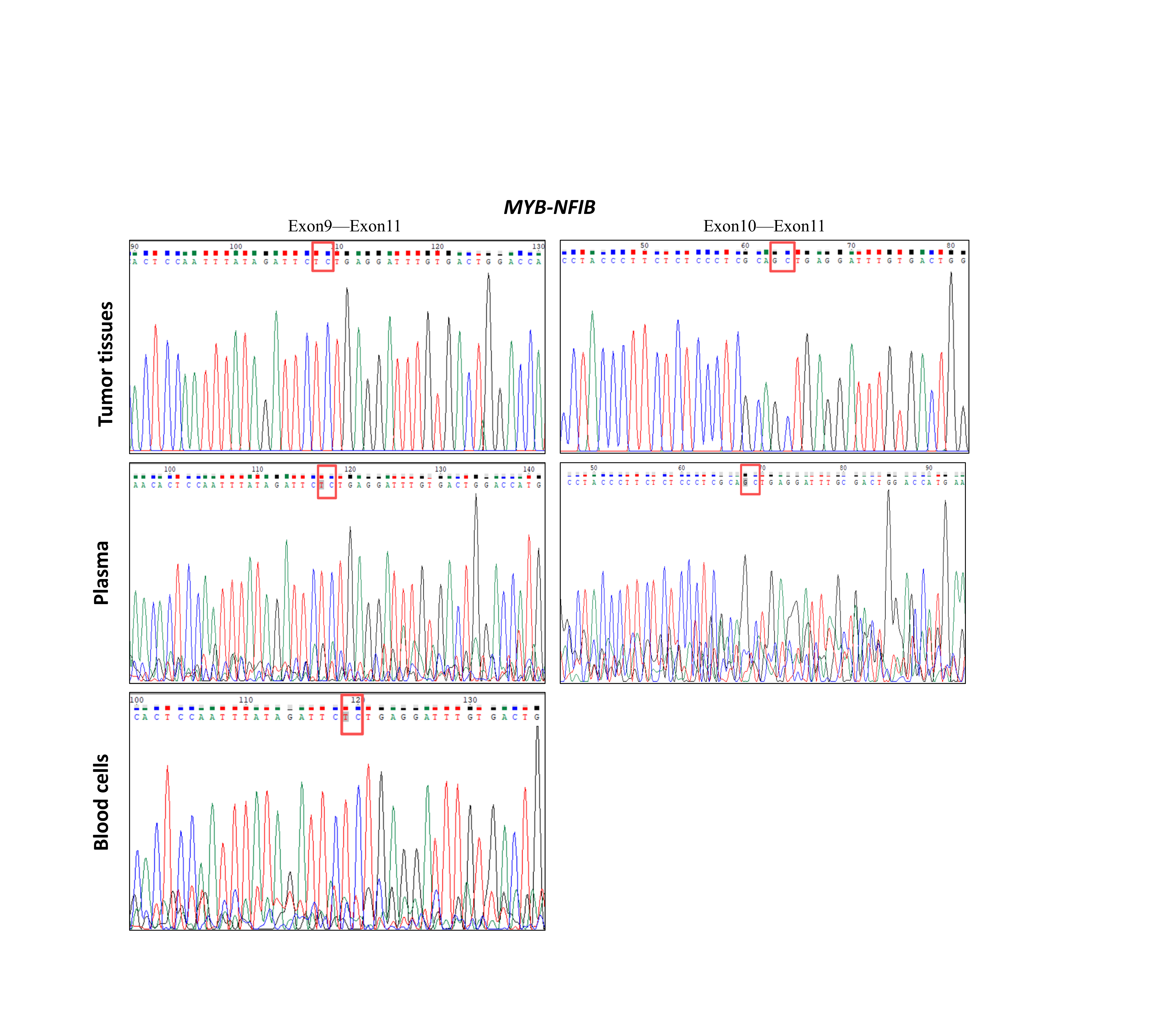

**Supplementary Fig S3. Tumor-derived *MYB‑NFIB* fusion gene detection in peripheral blood.** RNA samples extracted from tumor tissue and peripheral blood were analyzed by nested PCR followed by Sanger sequencing. Two fusion variants were identified in patient-matched primary **tumor tissue**: MYB exon 9–NFIB exon 11 and MYB exon 10–NFIB exon 11. Both fusion forms were detected in RNA from cell‑free **plasma**, whereas only one variant was detected in RNA isolated from **whole blood cells**.
